# Modelling live fuel moisture content at leaf and canopy scale under extreme drought using a lumped plant hydraulic model

**DOI:** 10.1101/2020.06.03.127167

**Authors:** N Martin-StPaul, J Ruffault, C Blackmann, H Cochard, M De Cáceres, S Delzon, JL Dupuy, H Fargeon, L Lamarque, M Moreno, R Parsell, F Pimont, JM Ourcival, J Torres-Ruiz, JM Limousin

## Abstract

- Water content in living vegetation (or live fuel moisture content, LFMC), is increasingly recognized as a key factor linked to vegetation mortality and wildfire ignition and spread. Most often, empirical indices are used as surrogates for direct LFMC measurements.
- In this paper, we explore the functional and ecophysiological drivers of LFMC during drought at the leaf and canopy scale using the SurEau-Ecos model, and a three years dataset of leaf and canopy scale measurements on a mature *Quercus ilex* forest, including an extreme drought. The model is based on forest hydrology and plant hydraulics and allows to simulate temporal variations of water potential and content at a daily time step. At leaf level, it simulates the relationship between water potential and water content by separating the apoplasm and the symplasm. Symplasm water content is modeled using the pressure volume curve theory, and apoplasm water content is modelled using the xylem vulnerability to cavitation. Fuel moisture content was upscaled to the canopy level by accounting for foliage mortality estimated from drought induced cavitation.
- The model was parameterized either with site-measured traits or using a calibration procedure, and compared with water potential and LFMC measured at leaf level, and NDVI variation measured at canopy level and taken as a surrogate for foliage mortality.
- At leaf level, LFMC prediction using measured hydraulic traits could be improved by considering year-to-year osmotic adjustments. At canopy level, foliage mortality due to drought induced cavitation was a key driver of LFMC decline during the most extreme drought.
- A sensitivity analysis showed that parameters driving soil water balance (leaf area index, soil water capacity, and regulation of transpiration) and parameters determining pressure volume curves are key traits driving LFMC dynamics at leaf level. At the canopy level, parameters that drives hydraulic failure were the most sensitive and included, both soil water balance parameters and hydraulic traits (the leaf vulnerability to cavitation) were the main drivers of LFMC decline during extreme drought.
- We also showed that under normal historic weather conditions, most variation of LFMC are linked to reversible symplasm dehydration, however under future, hotter and dryer conditions, most variations are due to the decline canopy of LFMC driven by foliage mortality.

## Introduction

Water content in vegetation is a key components of ecosystem responses to drought. Recent studies proposed it is likely to be related to drought induced mortality, a crucial outcomes of ongoing climate change (Martinez-Vilalta *et al.* 2019). One of the metric of water content or moisture content of plant is defined as the ratio of water mass to the dry mass. In the context of wildfire studies it is referred to as Live Fuel Moisture Content (LFMC), a variable that is is increasingly recognized as a critical factor in wildfire behavior (Rossa *et al.* 2016; Pimont *et al.* 2019a), hazard and activity (Dennison and Moritz 2009; Yebra *et al.* 2013; Ruffault, Curt, *et al.* 2018; Pimont *et al.* 2019b). Furthermore, being strongly determined by drought conditions, LFMC is one of the factors that might be responsible for observed and projected increase in wildfire hazard because of the increase in the frequency of drought event due to climate change.

Despite LFMC is important for our understanding of wildfire regime features and changes, in particular in foliage-fueled ecosystems, the precise role of LFMC is not well-established. A large part of these knowledge gaps is undoubtedly due to the fact that LFMC dynamic remain poorly understood and difficult to predict. Most studies that explore the drought-fire relationships do not consider explicitly the role of fuel moisture content in wildfire dynamics (Abatzoglou and Williams 2016; Turco *et al.* 2018; Ruffault *et al.* 2020). Instead, they rely on physical drivers of fuel moisture dynamics such as the vapor pressure deficit (VPD), or on empirical drought indices that are correlated with FMC, including dead FMC (Resco de Dios *et al.* 2015) and live FMC (Ruffault, Martin-StPaul, *et al.* 2018). Thus, approaches based on climate only account for climate, neglect sites and species physiological features, and hence, their performance for LFMC predictions is limited (Soler Martin *et al.* 2017; Ruffault, Martin-StPaul, *et al.* 2018).

In recent years, increasing attention has been paid to the understanding of the physiological drivers of live fuel moisture content, opening a field that has been called the “pyro-ecophysiology” by (Jolly and Johnson 2018), which should lead to the development of process-based approaches to predict fuel moisture content that should be valid under drought. This idea is supported by empirical findings showing the close relationships between LFMC and drought indices derived from functional approaches (Nolan *et al.* 2018; Ruffault, Martin-StPaul, *et al.* 2018; Pivovaroff *et al.* 2019). For instance, Ruffault *et al.* (2018) showed that soil water content was likely to be at least as good as empirical drought indices to predict LFMC dynamics. Others went further and showed that plant water potential is a good predictor of LFMC (Nolan *et al.* 2018; Pivovaroff *et al.* 2019).

Although water and carbon cycles in plants both dictate the variation of LFMC over the year (Jolly *et al.* 2014; Jolly and Johnson 2018), the mechanisms associated to the water cycle are the most influential drivers of live fuel moisture content dynamics over the course of a seasonal drought, whereas carbon cycles processes are particularly crucial during organ growth and development. Carbon-cycle processes influence LFMC through the phenological and physiological mechanisms that drive dry matter accumulation during growth, including photosynthesis, respiration and carbon allocation. Growth mostly occurs out of the dry season and, under drought conditions growth and photosynthesis generally ceased rapidly (Muller *et al.* 2011; Lempereur *et al.* 2015). It is thus reasonable to neglect these processes under extreme drought conditions in a first instance, and therefore assume that dry matter is constant during a strong drought.

Broadly speaking, water cycles processes involve different biophysical mechanisms that dictate water flows through the Soil Plant Atmosphere Continuum and within the plant. Mechanisms with a higher effect on LFMC during drought can be separated in two types. On the one hand, the forest hydrology processes that dictate how a given climate influences the soil water content and, therefore, the soil and plant water potential. This typically includes rainfall interception, infiltration and percolation, watershed hydrology, evaporation from the soil surface and plant transpiration. Leaf area of the stand and soil water-holding capacity, and the physiological processes involved in plant transpiration regulation (stomatal closure and minimum conductance, (Martin-StPaul *et al.* 2017; Cochard 2019; Duursma *et al.* 2019)) are key variables regulating these processes. On the other hand the plant dehydration mechanisms that dictate how plant dehydrates or desiccate at a given level of plant water potential. This later processes can be represented by separating the plant (or plant organ) into two main water reservoirs: the apoplasmic reservoirs (made of xylem conduits and call walls) and the symplasmic reservoir (made of living cells) (Tyree and Yang 1990; Martin-StPaul *et al.* 2017; Cochard *et al.* 2020). Each of them has a its own dynamics during decrease water potential that occurs during drought. For the symplasmic reservoir, water content dynamic is determined by the pressure volume curve theory, which states that water content dynamic during decreasing water potential essentially depends on cell wall elasticity and leaf osmotic potential (Tyree and Hammel 1972; Dreyer *et al.* 1990). In contrast, the water content dynamic of the apoplasm depends on the process of cavitation that leads to the change phase of water from liquid to gas (Tyree and Yang 1990; Hölttä *et al.* 2009; Martin-StPaul *et al.* 2017; Cochard *et al.* 2020). As water potential decreases and the amount of cavitation events in the xylem vessels increases, the amount of water in the apoplasmic tissue decreases.

At the canopy level, an additional effect of drought that influences canopy water content and that should be considered is the fact that leaves can progressively turn from live to dead under as a result of disconnection from the hydraulic system. Such foliage mortality would lead to a sharp drop in water content as dead foliage have very low moisture content values, between 5 and 20 % according to vapor pressure deficit and in the absence of rainfall (De Dios et al 2015). The processes that lead to foliage mortality could be linked to drought induced cavitation (Barigah *et al.* 2013; Urli *et al.* 2013) albeit this is a matter of research (Choat *et al.* 2018).

In this paper we explore the key parameters that drives fuel moisture content at the leaf and the canopy level during drought at the scale of a dry season. We used both a long term monitoring dataset at the leaf an canopy level and mechanistic model including the process described earlier.

The modelling approach includes both leaf level and canopy level water cycling processes, and bridges the gap between plant water balance model and plant hydraulics. It develops the links between plant water content and plant water potential by using the simplified version of the plant hydraulic model SurEau (Martin-StPaul *et al.* 2017; Cochard *et al.* 2020), and it uses a daily stand water balance model to predict soil water content by accounting for climate, soil properties and stand leaf area index (Rambal 1993; Granier *et al.* 1999; Ruffault *et al.* 2013; Cáceres *et al.* 2015).

The outputs of the model were tested on the experimental site of Puéchabon, a *Quercus ilex* coppice from which micro-meteorology, plant functioning, including leaf water potential and leaf water content data were collected and used to validate the model in details. We then propose a sensitivity analysis of the model to a few key traits and to climate change scenario to assess the potential impact of climate change on LFMC dynamics in Mediterranean forests.

## Materials and methods

### General approach

Our aim is to use a data-model analysis to explore the relevant processes that determines the dynamic of leaf and the canopy level moisture content under extreme drought. For this purpose, we present SurEau-Ecos, a model coupling a stand water balance model following the principle of previous models (Rambal 1993; Granier *et al.* 1999; Ruffault *et al.* 2013; Cáceres *et al.* 2015) and a simplified version of the soil-plant hydraulic model SurEau to simulate plant dehydration and desiccation (Martin-StPaul *et al.* 2017; Cochard *et al.* 2020). The stand water balance model is used to compute evapotranspiration, interception, soil water content, and soil water potential in the rooting zone. The SurEau model is used to compute plant water potential and live fuel moisture content at the leaf and canopy level as well as leaf mortality.

At leaf level we assumed that dominant processes include the relationships between plant water potential and water content, based on the theory of the pressure volume curves and for the symplasm and cavitation for the apoplasm. At canopy level we assume that the leaf mortality was caused by drought and could be related to cavitation. The main principles that dictate the dynamics of foliage desiccation are presented Figure 1. In the current stage, the model considers the desiccation of existing mature foliage and does not consider processes involved foliage growth and dry matter accumulation and carry over effect. Therefore, foliage and water transport capacity are re-initialised every year to their initial values.

**Figure 1:**
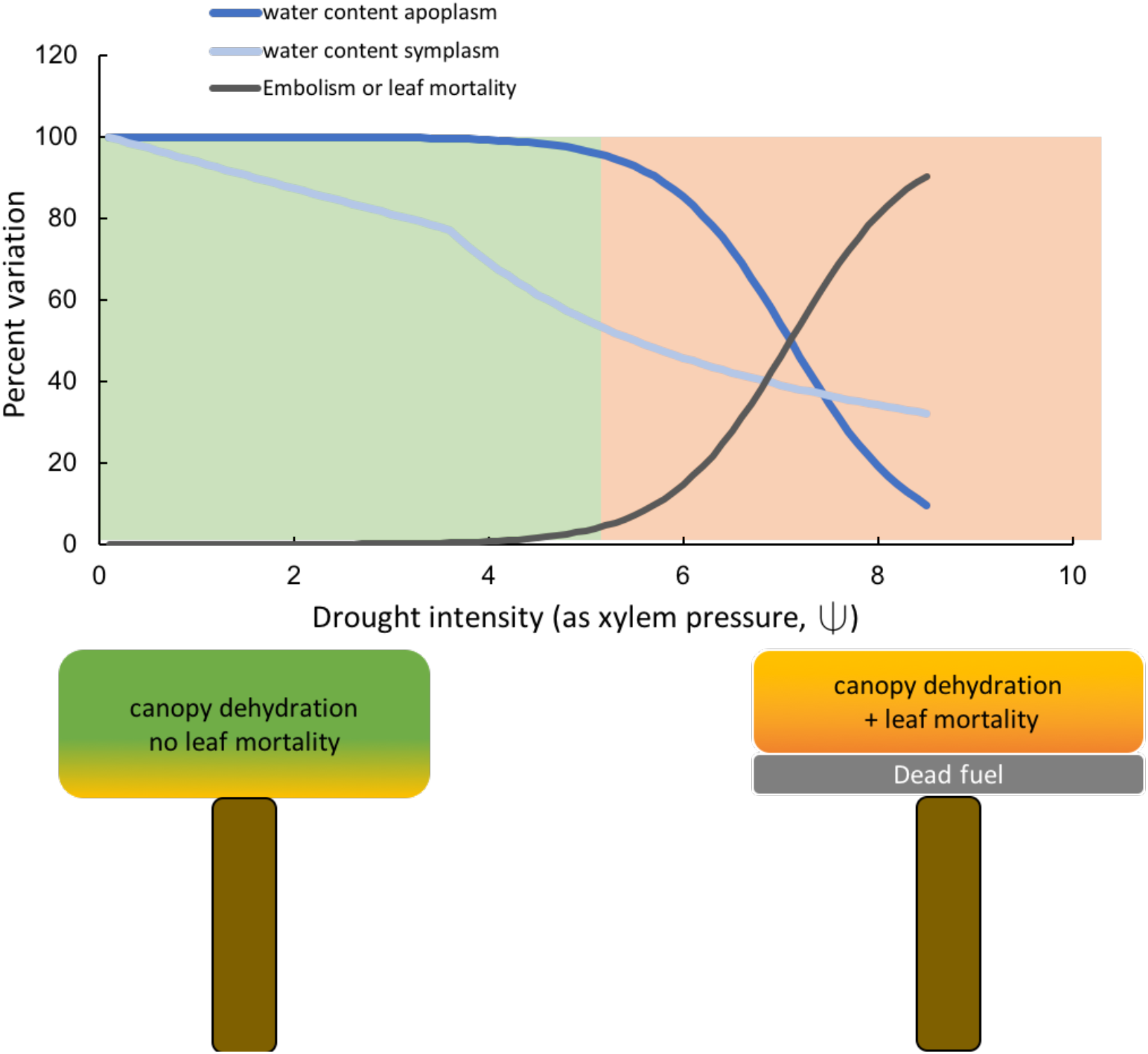
Schematic representation of the processes and traits involved in LFMC dynamic and used in the SurEau-Ecos model. Theoretical response of LFMC to plant water potential obtained thanks to a model describing the dynamic of two distinct compartment (the symplasmic compartment, living cells) and the apoplasmic compartment extracellular xylem water

In the following model descriptions, for clarity reasons, we separate the processes involved in (i) the water balance part of the model, (ii) the coupling between the plant hydraulic model and the water balance, and (iii) the computation of shoot level live fuel moisture content, the foliage mortality and the canopy level fuel moisture content. Afterwards, we present the data collected on the long term monitoring site of Puéchabon that are used to explore our underlying assumptions. We also carried out a sensitivity analysis of LFMC to ecological and physiological traits, as well as an exercise of stand canopy projections of LFMC under future climatic conditions.

### Model description

#### Overview of the water balance model and its coupling with the hydraulic model

The daily stand water balance model used in this study is based on equations previously developed for the SIERRA model (Mouillot *et al.* 2001; Ruffault *et al.* 2013), which follows the design principles of a previous water balance model (Rambal 1993; Granier *et al.* 1999; Cáceres *et al.* 2015). The detailed equations and principles having been published many times and provided into details in (https://vegmod.ctfc.cat/frames/medfatebook/), we only present the basic principles and the coupling with the hydraulic model SurEau. The model updates soil water content at a daily time step based on the water balance between precipitations (P) and water outputs:

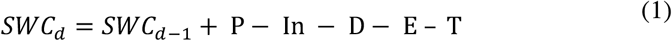

Where the amount of precipitation intercepted by the canopy (In), the soil evaporation (E), the transpiration of vegetation (T) and the drainage (D), are all expressed in *mm*. Soil is represented by a 3-layer bucket model which sizes can be parameterized to represent the available water capacity of the location of interest. Drainage from one layer to the other, and out of the system, occurs when the field capacity is overpassed. The interception of rainfall by the canopy is computed using the Gash equations (Gash *et al.* 1995). The soil evaporation is computed following (Mouillot *et al.* 2001), by considering soil surface potential evapotranspiration according to the Ritchie model (Ritchie 1972). Soil surface PET is obtained by dampening bulk air PET according to canopy leaf area index (LAI, m^2^ m^−2^) using an exponential decay (Mouillot *et al.* 2001). PET is computed using with the Priestley–Taylor equation as in (Ruffault et al 2013).

Plant canopy transpiration is computed by following the empirical bulk canopy limitation of the potential evapotranspiration (PET) proposed by (Granier *et al.* 1999). In addition, we introduce a stomatal regulation to drought through plant water potential (*ψ_plant_*), and a term of residual transpiration (*E_min_*, corresponding to losses through the leaf cuticle) computed as the product of the minimum conductance (*g_min_*) and the air vapor pressure deficit (VPD), assuming leaf and air temperature are equal. Both processes are key components of the dynamic of plant desiccation emphasized in this model. Averaged daily canopy transpiration rate (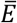, in mmol.m^−2^.s^−1^) then expresses as:

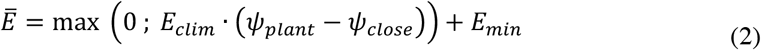

Where *ψ_close_* is a parameter corresponding to the water potential causing stomatal closure (taken as 95% of stomatal closure), *E_min_* is computed as the product of *g_min_* and vapor pressure deficit, and *E_clim_* is the boundary climatic transpiration, without stomatal regulation taken from (Granier *et al.* 1999):

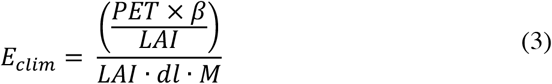

Where *M* the molar mass of water (kg.l^−1^) and *dl* day length (in seconds), and *β*:

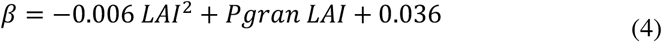

*P gran* is the Granier parameter that set the maximal transpiration per unit LAI that can be tuned if transpiration and LAI data are available.

The stomatal control on transpiration computed with (*ψ_plant_* − *ψ_close_*) is considered through the coupling with the plant hydraulic model SurEau. In this study, the preliminary version of SurEau (Martin-StPaul *et al.* 2017), originally dedicated to compute mortality due to “hydraulic failure” was used to compute plant water status, foliage water content and foliage mortality. As showed in the most recent developments, SurEau can also be applied to compute water quantity and desiccation under extreme drought (Cochard *et al.* 2020). In brief, it calculates soil-plant water transfer using water potential gradient and Fick’s law, and accounts for dynamic changes in hydraulic conductance with water potential decline due to drought induced cavitation. Then plant water content is computed by using the mechanistic links water potential to tissue water content. First, the Fick’s law can be applied to stand transpiration as:

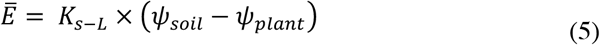

Where *E* is the average transpiration rate, *ψ_soil_* and *ψ_plant_* are soil and plant water potential. *K*_*S* − *L*_ is hydraulic conductance of the system (from soil to leaf). Soil water potential is computed from soil water content derived from the water balance model using the pedo-transfer function (Campbell 1974):

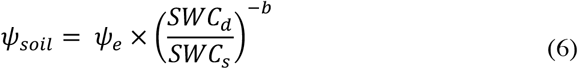

Where *ψ_e_* is the air entry water potential (MPa), *SWC_d_* and *SWC_s_* are the actual and saturation soil water content respectively (in mm as in Equation 1) and *b* is the power exponent of the soil moisture function. Note that this equation is generally expressed using fractional volumetric water content (*θ* in cm^3^.cm^−3^), as referred in Table 1. *K*_*S* − *L*_ has been simplified to consider the soil hydraulic conductance (*K*_*soil*_) and the plant hydraulic conductance (*K*_*plant*_) in series:

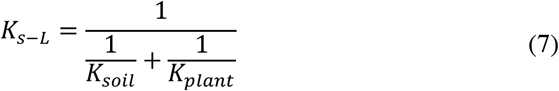

With (*K*_*soil*_) being computed by scaling the unsaturated conductivity equation of (Campbell 1974) with the Gardner-Cowan formulation (Gardner 1964; Cowan 1965) for accounting for the distance between soil and roots as in (Martin-StPaul et al 2017):

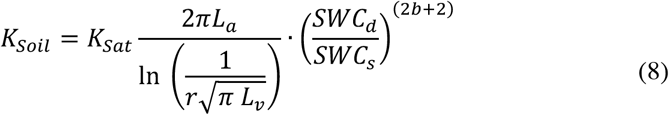

with *L_a_* and *L_v_* the fine root length per soil area and soil volume, respectively and *r* the fine root radius.

Equaling 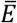 from equation 2 and 5 because of water mass balance and rearranging terms leads to the following solution for *ψ_plant_*:

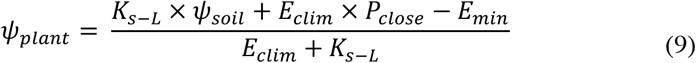

**Tableau 1:**
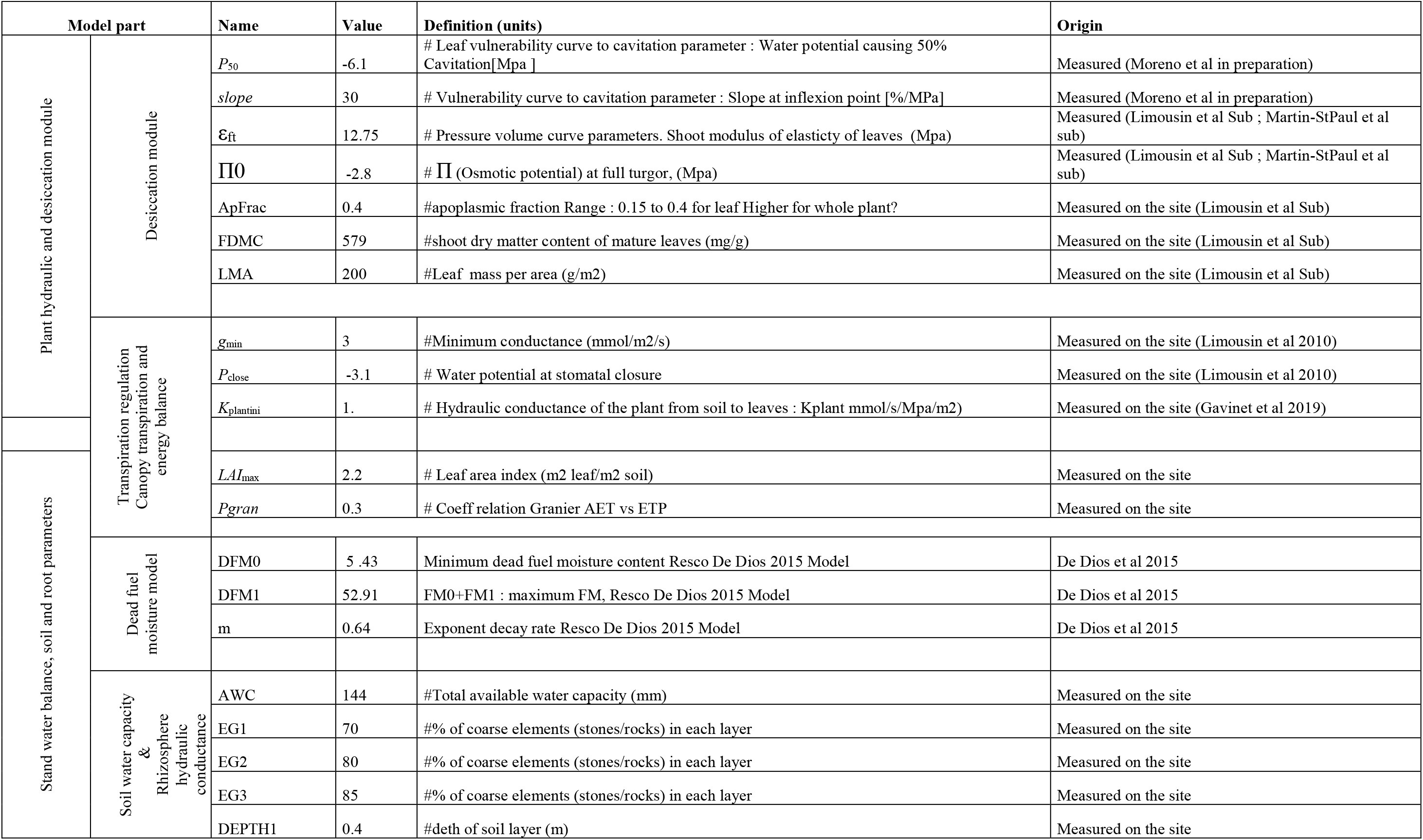
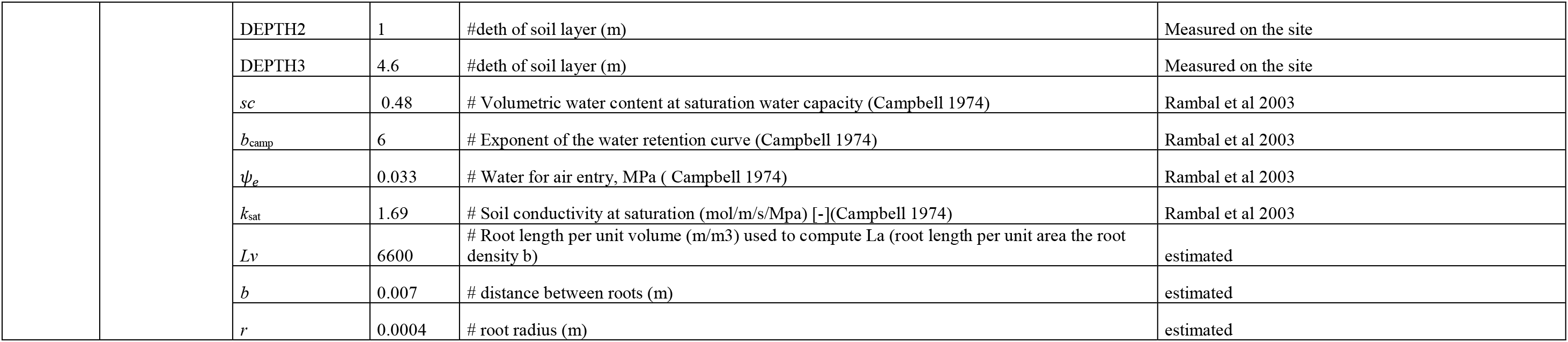
Model default parameters for *Quercus ilex* stand at Puéchabon Forest. These parameters were mostly measured on the sites. Note that parameters Π0 and Esymp were also adjusted using the temporal dynamics of leaf LFMC and water potential (see Table 2).

However, when this solution leads to *E*_*clim*_ · (*ψ_plant_* − *ψ_close_*) < 0, corresponding to full stomatal closure, only the cuticular losses drive the flux and *ψ_plant_* write:

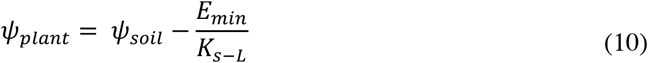

In practice, equations are solved as follows. *E*_*clim*_ is first computed from equation 3 and the day length (*i.e.* corresponding to the transpiration rate without considering stomatal closure), *ψ_soil_* and *K*_*S* − *L*_ are taken from the previous time step, so that *ψ_plant_* can be computed using equation 9 and reinjected into equation 2 to compute the plant transpiration *TR.* At the end of the procedure, *ψ_plant_* is used to compute the loss of hydraulic conductance due to cavitation as well as dehydration involved in LFMC computation (see next section).

Along a dry season, *K*_plant_ can decrease according to level of drought induced cavitation (Cruiziat *et al.* 2002). This process corresponds to a phase change of water from liquid to gas that lead to an air embolism in the xylem which precludes sap circulation and empty the vessels from water. *K_plant_* is computed as a function of initial (pre-drought) plant conductance (*K_plantini_*) and the fractional loss of conductance (*LC*) due to drought induced embolism:

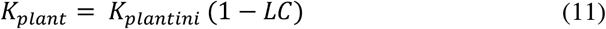

The fractional loss of conductance of the plant xylem is assumed proportional to the level of drought-induced *Embolism* in the xylem. It occurs when water potential drops below the intrinsic capacity of the plant xylem to support negative water potential. *Embolism* is computed by using the sigmoidal function (Pammenter and Vander Willigen 1998):

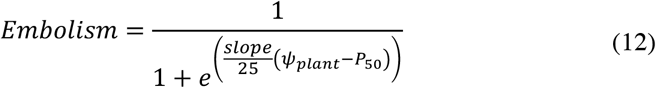

Where *P*_50_ (MPa) is the water potential causing 50% loss of plant hydraulic conductivity and *Slope* (%/MPa) is a shape parameter describing the rate of embolism spread per unit water potential drop at *P*_50_. To simplify the resolution of water potential and transpiration regulation, LC assumed to be constant during a time step. *K_plantini_* is reinitialized to its maximum value at the beginning of each year, assuming that tissue growth reestablished the transport capacity of the plant.

#### Modelling Live fuel moisture content at the leaf and canopy scale

##### a Live fuel moisture content at the shoot level

The water content at the shoot level is computed from static relationships between water potential and water content (Martin-StPaul *et al.* 2017; Cochard *et al.* 2020). The model considers that foliage has two main different water reservoirs -apoplasmic and symplasmic (Tyree and Yang 1990), each of them having its proper response to *ψ_plant_*.

The relative water content of the symplasmic compartment (*RWC_s_*), decrease as *Ψ_plant_* becomes more negative according to the well-known pressure volume curves equations (Tyree and Hammel 1972). PV curves depends on two main parameters, the osmotic potential at full turgor (π_0_, MPa) and the elasticity of cell walls (ε % / MPa). Importantly these parameters can be easily obtained from laboratory measurement datasets, available for many species (Bartlett *et al.* 2012; Martin-StPaul *et al.* 2017). *RWC_s_* is the minimum values of the two possible solutions, when the leaf is still turgid (Ψ > Ψ_*tlp*_, the turgor loss point):

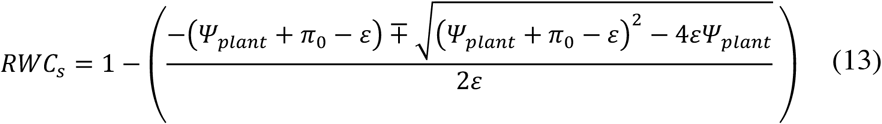

Alternatively, when *Ψ_plant_* ≤ *Ψ_tlp_* (if water potential has overpassed the turgor loss point), the solution is:

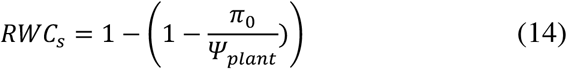

Then, the relative water content of the apoplasmic compartment (i.e. extracellular water contained in the xylem,), declines in response to plant water potential (*Ψ_plant_*) according to the level of embolism. It is obtained from the vulnerability curve to cavitation (Equation 11) that can estimated for the organs of different species through the percent loss of conductivity in response to water potential (Choat et al 2012; Martin-StPaul et al 2017). We assume that *RWC_a_* increases proportionately with the rate of embolism and consider that embolism is not reversible at the scale of a dry season:

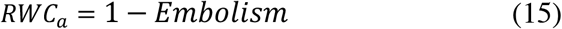

To obtain the live fuel moisture content on a dry mass basis, we expressed *LFMC* as the sum of the water content in the symplasm and the apoplasm relative to the dry mass weighted by they respective volumetric fraction:

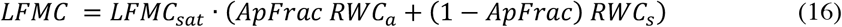

With *ApFrac* the fraction of apoplasmic tissue of the shoot (that can be derived from pressure volume curves) and *LFMC_sat_* the live shoot moisture content at saturation (in *g* H_2_O. g^−1^). For an easier parameterization, *LFMC_sat_* can be derived from more classical traits such as *LDMC* or leaf succulence (*S,* in *g* H_2_O.m^−2^) and leaf mass per area (*LMA*, in *g* dry matter .m^−2^).

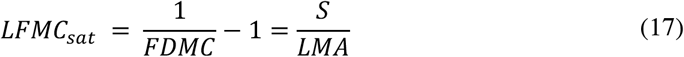

Equations 12 to 17 allow to link water content and water potential and can be applied to field empirical measurement, or by using pressure volume curves and vulnerability to cavitation (see below). Note that we assume that the “bulk plant water potential” modelled is the same in the apoplasmic and the symplasmic compartments. In order to compare model output with field data (see below), we used the simulated soil water potential (Ψ_*soil*_) as predawn water potential and Ψ_*plant*_ as a midday water potential.

##### b Canopy scale fuel moisture content and foliage mortality

The total canopy scale fuel moisture (or total canopy water load, *TFM_can_*, gH2O.m^−2^) can be computed by summing the amount of water in live and dead foliage:

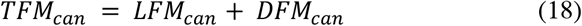

Where *LFM_can_* is the live canopy scale fuel moisture (or live canopy water load, g H_2_O.m^−2^) and *DFM_can_* is the dead canopy scale fuel moisture (or dead canopy water load, g H_2_O.m^−2^). *LFM_can_* is computed daily by scaling leaf level live fuel moisture content to the canopy dry mass by using the leaf area index (*LAI*) and the leaf mass per area (*LMA*).

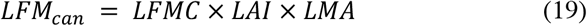

Whereas *LMA* is assumed constant (as growth process and dry mass accumulation are ignored); *LAI* can decrease during a dry season due to foliage mortality in case of extreme drought. In order to account for this phenomenon, we included a leaf mortality curve in the model to update living *LAI* and to compute the dead foliage. This phenomenon also induced a decrease in moisture content of the total canopy as dead leaf moisture content can reach values as low as 5 to 10 % during dry conditions and follows a dynamic imposed by air vapor pressure deficit (Resco de Dios *et al.* 2015).

To model leaf mortality, we made the parsimonious assumption that drought induced embolism leads to a proportional increase in leaf mortality, in agreement with many previous studies on forest trees showing a concordance between canopy mortality and hydraulic failure (Blackman *et al.* 2010; Barigah *et al.* 2013; Urli *et al.* 2013; Li *et al.* 2015). In practice, leaf mortality is computed at each time step as a fraction of the maximum leaf area index and the variation of embolism between the current and the previous time step:

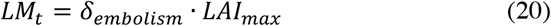

Where *LM_t_* (in m^2^.m^−2^) is the leaf mortality computed at each time step, *LAI_max_* is the maximum yearly value of *LAI* and *δ*_*embolism*_ is the variation of embolism between two consecutive time steps (i.e. during one day). Because a native level of embolism on the order of 5% is frequently recorded in this species (Martin-StPaul *et al.* 2014), the effect of embolism on leaf mortality is not accounted below this threshold. The *LM_t_* is then used to update the actual leaf area index and the dead leaf area compartment:

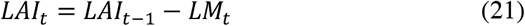

With *LAI*_t_ and *LAI*_t-1_, the *LAI* of the current and previous day respectively. The dynamic of dead leaf area index (*DLAI*) is also computed as:

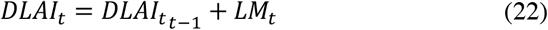

With *DLAI*_t_ and *DLAI*_t-1_, the dead *LAI* of the current and previous day respectively. Then the fraction of moisture of dead fuel (*DFMC*, % of dry mass) can be modelled as a function of vapor pressure deficit (VPD) with the semi-empiric model of Resco De Dios et al (2015), and used to compute the total dead fuel moisture content of canopy assuming that dead leaves stay on the tree.

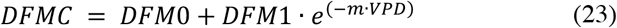

Where *DFM_0_*, *DFM*_1_ and *m* that can be adjusted to empirical data or taken from Resco De Dios et al (2015). Knowing *DFMC* and *DLAI*_t_, the dead fuel moisture of the canopy (*DFMC_can_*) is computed:

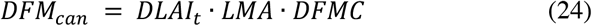

### Model application to a *Quercus ilex* canopy at the Puéchabon site

The model was applied on the *Quercus ilex* forest stand of Puéchabon. The Puéchabon site is located 35 km north-west of Montpellier (southern France), on a flat plateau, in the Puéchabon State Forest (3°35’45”E, 43°44’29”N, 270 m ASL). The forest has been managed as a coppice for centuries and the last clear cut was performed in 1942. In 2011, the top canopy height was 5.5 m on average. The stem density of *Quercus ilex* evaluated on 4 plots larger than 100 m^2^ was 4703 (± 700) stems ha^−1^. The climate is Mediterranean with a mean annual precipitation of 903 mm and a mean annual temperature of 13°C (on average 1984-2011). The very shallow bed rock imposes a strong constraint on water availability: the volumetric fractional content of stones and rocks averages 0.75 for the top 0-50 cm and 0.90 below. The vegetation is largely dominated by a dense overstory of the evergreen oak *Quercus ilex L.*, so that the application of our stand scale monospecific model is consistent. The site has historically been used to test functional models on forest stand in the Mediterranean (Reichstein *et al.* 2002; Davi *et al.* 2006; Keenan *et al.* 2011; Ruffault *et al.* 2013).

Studied years spanned from 2016 to 2018, and included one of the most extreme drought recorded at the site (year 2017) that caused partial mortality and desiccation of the canopy. During the summer drought (May to October) of the studied period, the water potential and leaf fuel moisture content were measured at predawn and at midday on five trees at approximatively 3 weeks intervals. Two or three leafy shoots per tree were sampled from the upper part of the canopy, sealed in a plastic bag in a cooler, and their water potential was measured within 2 hours using a Scholander pressure bomb (PMS1000, Corvallis, Oregon, USA). Three to six leaves were collected concomitantly from the same trees for LFMC measurements, stored in sealed plastic bags right upon collection and kept in the cooler. At the laboratory, leaf samples were weighted fresh, and then dried an oven at 60°C for 48 hours and weighted again to compute LFMC. Leaf LFMC measurements on mature current year leaves only were used. *Quercus ilex* bud burst generally occurs at the end of March and the new shoots expand and maturate until early July. Consequently, at the beginning of each measurement season (May to June), it happened that only previous year leaves were available or that current year leaves were not fully mature which were not used for this study. Daily meteorological variables used to feed the model include rainfall, temperature, radiation, vapor pressure deficit, wind speed and were taken from the site meteorological station. Additional ecophysiological data used here also include sap flow measurements, that have been continuously been recorded since 2003 (Limousin *et al.* 2009; Gavinet *et al.* 2019).

#### Parameterization of the water balance model

Model parameters for the water balance part were taken from (Rambal *et al.* 2003; Ruffault *et al.* 2013) or adjusted with more recently available *in situ* measurements (Table 1). The leaf area index was initialized at a maximum value *LAI*_max_ of 2.2. The total available water capacity was estimated at 145 mm by using eddy-flux measurements. The maximum soil depth of the soil layer of the water balance model was estimated to 4.6 m by taking into account the soil volumetric rock fraction. This value was used to calibrate the water balance model. The total hydraulic conductance was set using sap flow and water potential gradient (between predawn and midday) measurements (Gavinet *et al.* 2019). The water potential causing stomatal closure (*P_close_*) was set to −3.1 MPa based on concurrent measurements of leaf gas exchange measurements and water potential (Limousin *et al.* 2009; Martin-StPaul *et al.* 2012). Minimum leaf conductance (*g_min_*) is considered here as a water balance parameters as it determines (along with *VPD*) the rate of water loss when stomata are close. It was estimated with data from pressure volume curves measurements (see below). The water balance component of the model has been explored previously, to complement prior validation for the studied years, we compared model outputs with *in situ* measurement of predawn and midday water potentials and sap flow.

#### Model evaluation at the shoot and stand level

In order to decompose the drivers of LFMC at the leaf level we first applied only leaf level LFMC equations (Equation 12 to 17) to field measurements of water potential and LFMC, and then evaluated the predictability of LFMC of the full coupled model.

Two main types of hydraulic traits are needed to parameterize the set of equations that determines the relationship between water potential and live fuel moisture content (equation 12 to 17): the leaf pressure volume curves (leaf PV curve) and leaf vulnerability to cavitation (leaf VC). Such measurements were performed during 2018 and 2019, on six trees of the site, and the detailed description of these data are provided in (Limousin et al in prep; Moreno et al in prep). In brief, pressure volume curves were measured using the bench dehydration technique (Dreyer *et al.* 1990) during the spring and summer 2018 on mature leafy shoots (10 to 15 cm length). The parameters include the osmotic potential at full turgor (π_0_) and the modulus of elasticity but also the fuel dry matter content (*FDMC*), the apoplasmic fraction (*ApFrac*). Note also that the minimal conductance (*g_min_*, that serve in the water balance model, see above) is derived from the last portion of the pressure volume curves. Among these traits some can exhibit some degree of plasticity to drought (Bartlett *et al.* 2012). This is particularly the case of “osmotic adjustment”, which corresponds to an increase of π_0_ with increasing drought, and that has been found to occur in response to rainfall exclusion in *Quercus ilex* at the Puéchabon and Font-Blanche sites (Limousin et al in preperation). As PV curves were measured only at one date (in march 2018) on shoots that included two generation of leaves (2016 and 2017), we cannot identify if osmotic adjustment has occurred between years and modified the relationship between water potential and LFMC. We therefore also used the LFMC and water potential data monitored on the site along the 3 measurement years to calibrate these parameters through parameter optimization (see the section below *Calibration*).

Early in 2019, leaf vulnerability to cavitation was assessed using the optical technique (Brodribb *et al.* 2016) applied to cut branches from the same six trees of the site using similar protocol to (Lamarque *et al.* 2018). In brief, large branches (>1m) were cut in the field, their cut bases immerged the water and brought to the laboratory. Once at the laboratory, the branches were placed on a bench with two leaves to three leaves scanned at a regular interval of 5 minutes using a flat scanner. One stem psychrometer (StemPsy1, ICT International, Armidale, NSW, Australia) was placed on the branch and stem water potential was recorded every 30 min throughout branch desiccation. The accuracy of psychrometer readings was cross-validated three to four times a day by *Ψ*_leaf_ measurements on adjacent leaves that had been covered for at least 2 h with aluminum foil and wrapped in a plastic bag using a Scholander pressure bomb.

##### a Leaf level LFMC model evaluation

We first evaluated the *LFMC* prediction at leaf level by using equations 12 to 17, that relates water potential and leaf moisture content by using the leaf monitoring on the Puéchabon site. To do so, the set of equation was used to predict predawn and midday LFMC data (monitored from 2016 to 2018) from predawn and midday water potential data. Three different parameterizations were tested. First we parameterized the equations by using measured values derived from pressure volume curves and vulnerability curves to cavitation (see above). Second, we proceed to an overall parameter optimization by using the whole dataset of LFMC and water potential monitoring. Finally, to account for the possible plasticity of pressure volume curves traits (in particular π_0_) from year to year, we proceed to a year by year parameter optimization by considering the data from each year independently. Note that we limited parameters optimization to pressure volume curves parameters (π_0_, 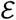, *FDMC*) that are known to adjust according to drought (Bartlett et al 2012; Limousin et al in preparation). Conversely, the cavitation vulnerability curve was not adjusted because cavitation occurs mostly outside of the range of water potential data measured in the field and is expected to exhibit little changes in response to drought variations (Limousin *et al.* 2010; Martin-Stpaul *et al.* 2013).

We also applied the coupled SurEau-Ecos during the three studied years by using the best parameterization found from Equation 12 to 17. We evaluated the ability of the model to simulate stand level foliage mortality recorded by using Normalized Difference Vegetation Index (NDVI) measurements made using a sensor positioned above the canopy. We computed an index of foliage change during the summer drought as the relative variation in NDVI between leaf maturity (around early July) and the end of the summer.

#### Sensitivity analysis and applications

##### a. Evaluation of traits responsible for leaf and canopy desiccation under current and future climate

Finally, we analyzed the sensitivity of leaf and canopy moisture content to different traits and parameters that are expected to be relevant to define desiccation. We selected a group of parameters that differ according to the type of processes in action: On the one hand, we computed the sensitivity to leaf and hydraulic traits that define the relationship between leaf level moisture content and water potential -- including the pressure volume curve parameters (π_0_, 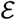) and the vulnerability to cavitation parameters (*P*_50_, *Slope*) -- and on the other hand we quantified the sensitivity of the model to parameters that affect the water balance of the stand -- including the water potential causing stomatal closure (*P*_close_), the minimum leaf conductance (g_min_), the leaf area index (LAI) and soil water capacity (*SWC*). For each parameter we explored how a variation of 40 % around its initial value (i.e. from −20 % from the initial to +20% from the initial value, Table 1) affects the minimum LFMC at the leaf and at the canopy level and the foliage mortality. This exercise was performed by simulating these variables for the period 1999-2019 and extracting the 5^th^ quantile of LFMC at leaf and canopy level and the 95^th^ quantile of leaf mortality.

In order to explore the LFMC sensitivity to climate change we performed simulations with the reference parameters for the period 2015-2100 using climate projection from one GCM-RCM (MPI-RCA4) under the RCP8.5 scenario. These climate simulations have been chosen to represent the average climate trajectory among 5 contrasted GCM-RCM according to Fargeon et al (2020). For this application, we used default plant parameters (Table 1). Some key parameters were re-initialized each year assuming that stand remain identical in terms of species and structure, that is we imposed a full recovery of plant conductance and leaf area index and no species turn-over. This should be considered as a theoretical exercise and not as a projection. We extracted yearly minimum LFMC values at leaf and canopy level and foliage mortality.

## Results

### Leaf level fuel moisture content dynamic relation to plant water potential

Figure 2 shows the relationship between live leaf moisture content and water potential measured and modelled (Equation 12 to 16) for each measurement year by using three different calibrations: (1) calibration with measured parameters, (2) calibration with fitted parameters using all data LFMC and water potential data together (i.e. 2016-2018 monitoring) and (3) calibration by fitting parameters on each year independently. The later procedure assumes that pressure volume curves adjustment can affect the relationship between LFMC and water potential from year to year. Parameters used for the calibration (either measured or optimize) as well as goodness of fit for the predictability of LFMC are reported on Table 2. We observed that LFMC simulated with parameters measured on the site (pressure volume curve equations) shows relatively good agreement on average with the data (RMSE = 5.2, R2=0.55, Table). However, for low water potential values (<−3.1 MPa), we noticed an underestimation of LFMC for year 2016 and an overestimation for 2017 and 2018. By proceeding to a calibration with the full data set at once, we obtained parameters very close from the parameters measured using PV curves (Table 2), and these values conducted to slightly improve the predictability of LFMC (RMSE=4.6, R2=0.59), but, as for calibration with measured parameters, LFMC was underestimated in 2016 and overestimated in 2017 and 2018. By optimizing parameters for each year independently, we found that optimal parameters vary and this leads to a substantial improvement of the overall predictability of LFMC (RMSE=3.7, R2=0.73). In particular, the overestimation found in 2016 and the underestimation found in 2017 and 2018 disappeared. Changes in parameter values particularly affected π_0_ that increased with the drought intensity of year (i.e. minimum value of water potential), from −2.5 year 2016 to years −3.2 2017 and 2018

**Table 2:** parameters values (i.e. traits) and goodness of fit used for the calibration of the LFMC prediction at leaf level (Equations 12 to 17, main text)

|  |  | Traits |  |  | goodness of fit<br>leaf level model |  | goodness of fit<br>coupled model |  |
| --- | --- | --- | --- | --- | --- | --- | --- | --- |
| | | $\Pi_0$ | $\varepsilon_{ft}$ | $FDMC$ | RMSE | R2 | RMSE | R2 |
| Calibration type |  | (MPa) | (MPa) | (mg.g<sup>-1</sup>) | % | - | % | - |
| Measured parameters | - | -2,81 | 12,75 | 579 | 5.22 | 0.55 | 7.5 | 0.3 |
| Fitted parameters | All years | -2,67 | 15 | 567 | 4.63 | 0.59 | 7 | 0.3 |
| Fitted parameters (per year) | 2016 | -2,45 | 15 | 573 | 3.67 | 0.73 | 5.3 | 0.56 |
|  | 2017 | -2,94 | 7 | 554 |  |  |  |  |
|  | 2018 | -3 | 15 | 566 |  |  |  |  |

**Figure 2:**
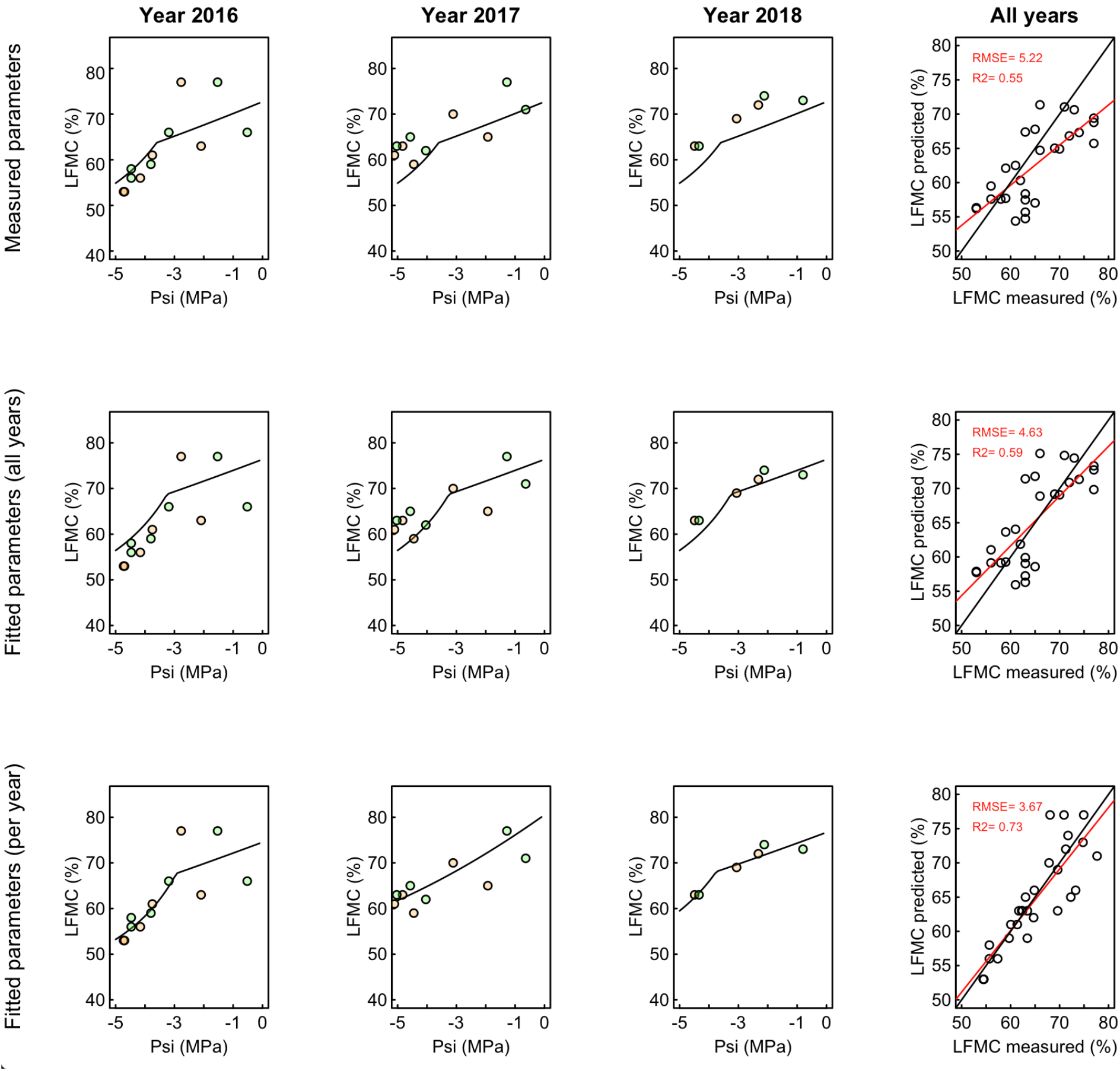
Leaf level module of moisture content and the role of osmotic adjustments. Measured and modelled relationships between leaf live moisture content (LFMC, %) and leaf water potential (***ψ***, MPa) for the three studied years at the Puéchabon site. Measurments at predawn and midday are shown in green and orange respectively. In the upper panels measured parameters were used to model LFMC as a function of ***ψ***, (Equations x to y). In the middle panels parameters were calibrated using field LFMC and water potential data by considering all the data together. In the bottom panels parameters were calibrated using field LFMC and water potential data by considering each year independantly. RMSE and R^2^ of the fitted versus measured LFMC values are indicated, parameters and goodness of fit are also are reported in table 2.

### Evaluation of SurEau-Ecos coupling the water balance and desiccation model

#### Water balance

The water balance part of the model has previously been validated at the site against sapflow data and water potential (Ruffault et al 2013). Here we present an additional validation of predawn water potential data for the three years studied (Figure 3). Model simulations show a good agreement with measurements with an overall R^2^ =0.91 for midday water potential and 0.89 for predawn water potential) and an overall RMSE = 0.5 MPa (0.4 for midday water potential and 0.6 for predawn water potential).

**Figure 3:**
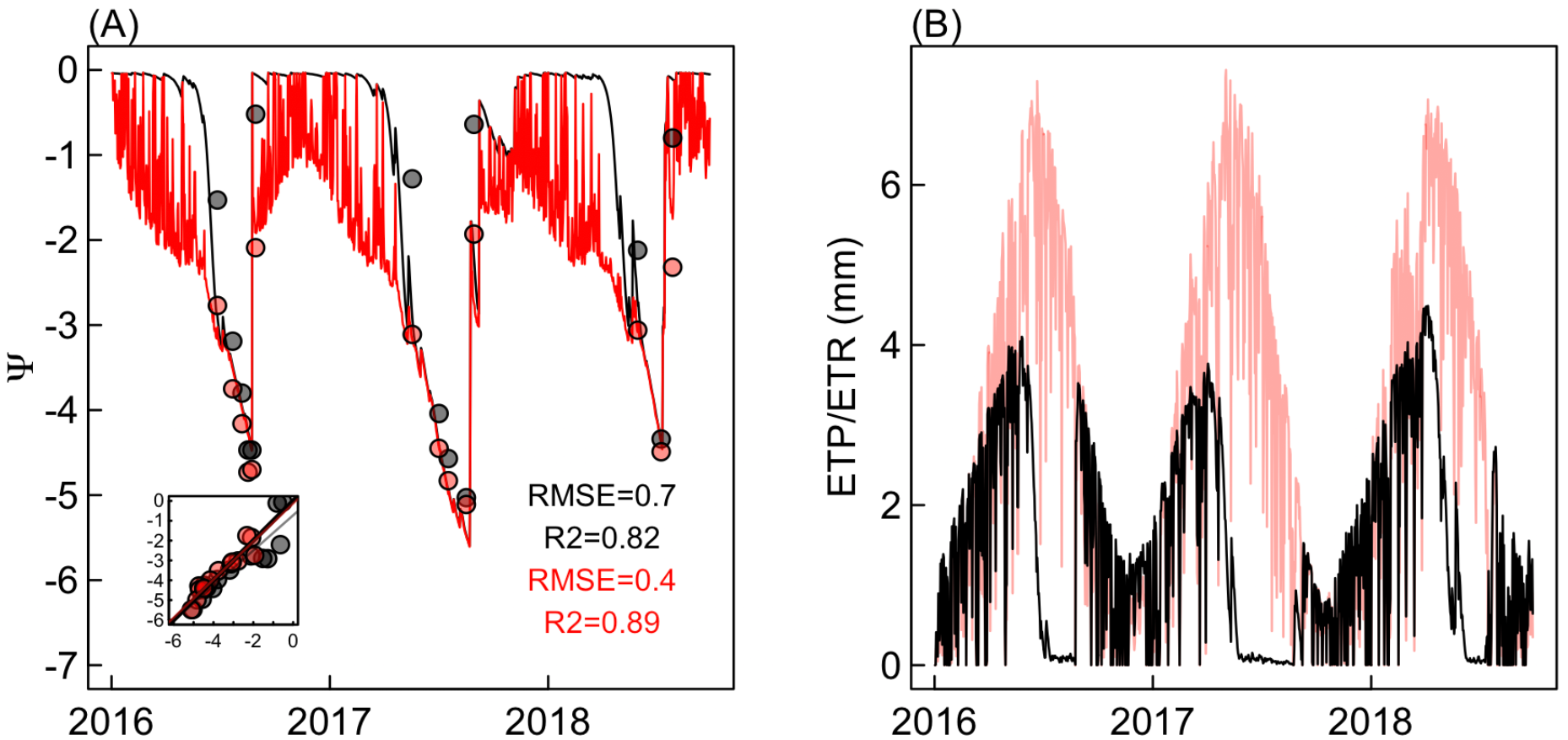
Water balance evaluation of SurEau-Ecos at Puéchabon.

#### Live fuel moisture content at leaf scale

An evaluation of the ability of the coupled model to predict the dynamic of leaf live fuel moisture content under the same calibration procedure as for leaf level evaluation is presented on Figure 4 and Table 2. In general, the performance of the coupled model to predict LFMC was lower than at the leaf level evaluation, but the same ranking between calibration procedure was found (Table 2). By using measured parameters, the RMSE and R^2^ were 7% and 0.3 respectively with pronounced underestimation bias during the peak of the drought in 2017 and 2018. Only marginal improvement was obtained by using parameters calibrated using all LFMC and water potential data (R^2^=.3 RMSE=7%). However by using parameters adjusted per year we obtained a significant improvement over the overall predictability of LFMC (R^2^=0.56, RMSE=4%) which was associated to a reduction of the overestimation bias during the peaks of dry years 2017 and 2018 (Figure 4). For the last set of parameters we presented the dynamic of leaf level water content in the symplasm and in the apoplasm (Figure 4d). This shows that most of the decrease in LFMC during drought is associated with symplasmic water decrease, whereas apoplasmic decline due to drought induced cavitation only slightly changed during the extreme 2017 drought.

**Figure 4:**
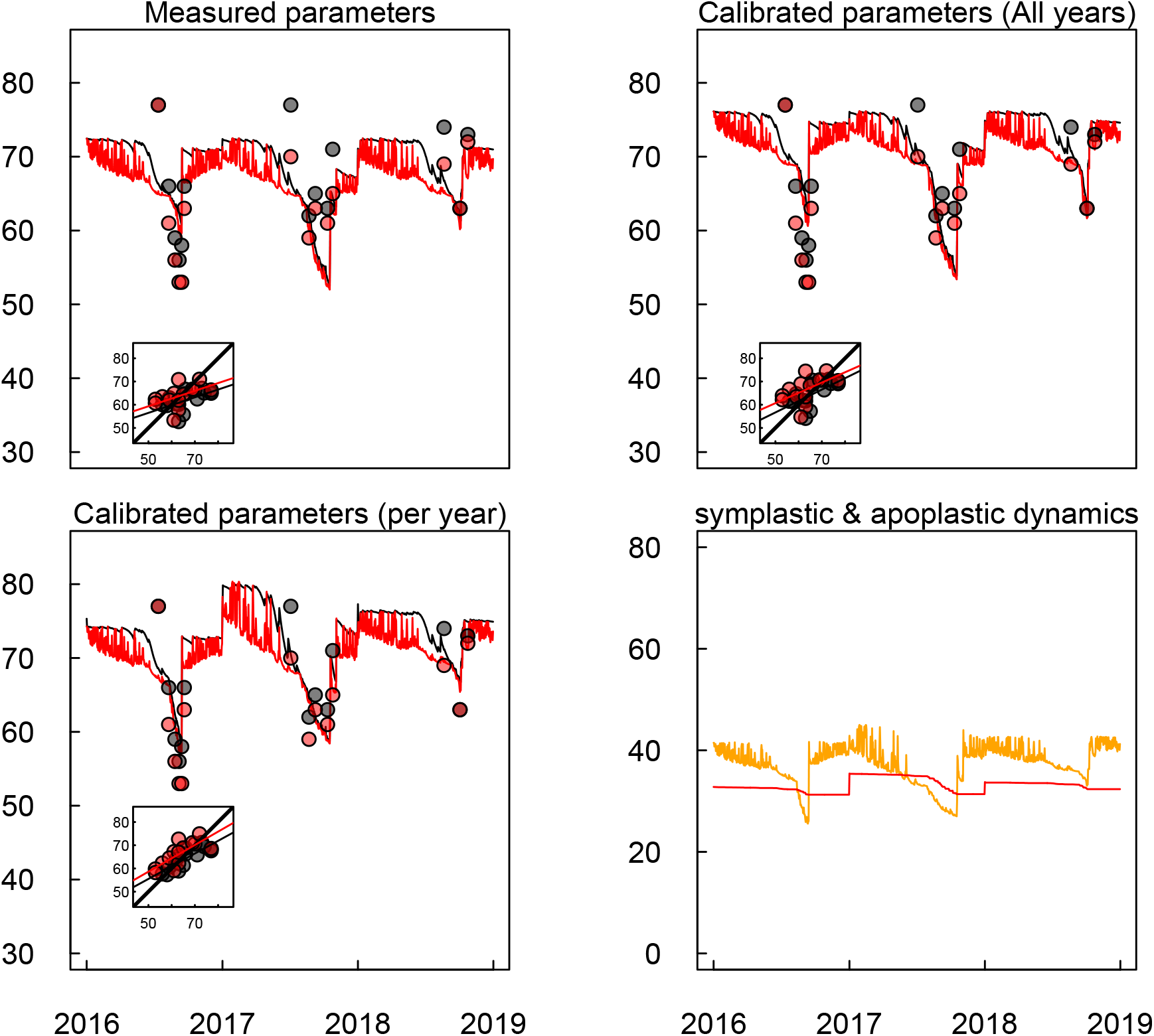
Leaf level dynamic of fuel moisture content for the three studied years modelled with the coupled SurEau-Ecos with different parameterization and measured. Values at predawn and midday are shown in black and and red respectively. The different parameterization were used for the equations describing LFMC as a function of ***ψ*** (Equations 12 to 17), leaf level equations (Figure 2, Table 2). On the first panel, traits measured on branches were used. In the second panel, parameters were calibrated using field LFMC and water potential monitoring by considering all the data together. In the third panel, parameters were calibrated using field LFMC and water potential monitoring by considering each year independantly. RMSE and R2 of the fitted versus measured LFMC (showed in the insert) values are reported in Table 2. The last panel shows the symplasmic (orange) and apoplasmic (red) moisture content dynamics simulated using the third parameterization for the three studied years. ***c***

#### Live fuel moisture content at canopy scale

Canopy water content dynamics is computed by integrating changes of living leaf water content and dead leaf water content and by upscaling them at the canopy level by accounting for possible leaf mortality. Accordingly, the canopy water content dynamics sharp decreases during summer drought in accordance with leaf level dynamics (Figure 5a). During the extreme drought 2017 (and to a lesser extent in year 2016) we observed that fuel moisture content drop was due to leaf mortality. We recall, that leaf mortality is computed proportionally to embolism and its water content is then computed as a function of vapor pressure deficit following (De Dios et al 2015). The increase in leaf mortality in 2017 reached almost 20% in accordance with embolism increase. The greater defoliation simulated during 2017 compared to 2016 and 2018 was consistent with the relative change in *NDVI* during the summer season (Figure 5c), as well as we the observation of leaf browning on the made on field this year (Figure 5d) and the observation of significant cavitation rate this year (data not shown).

**Figure 5:**
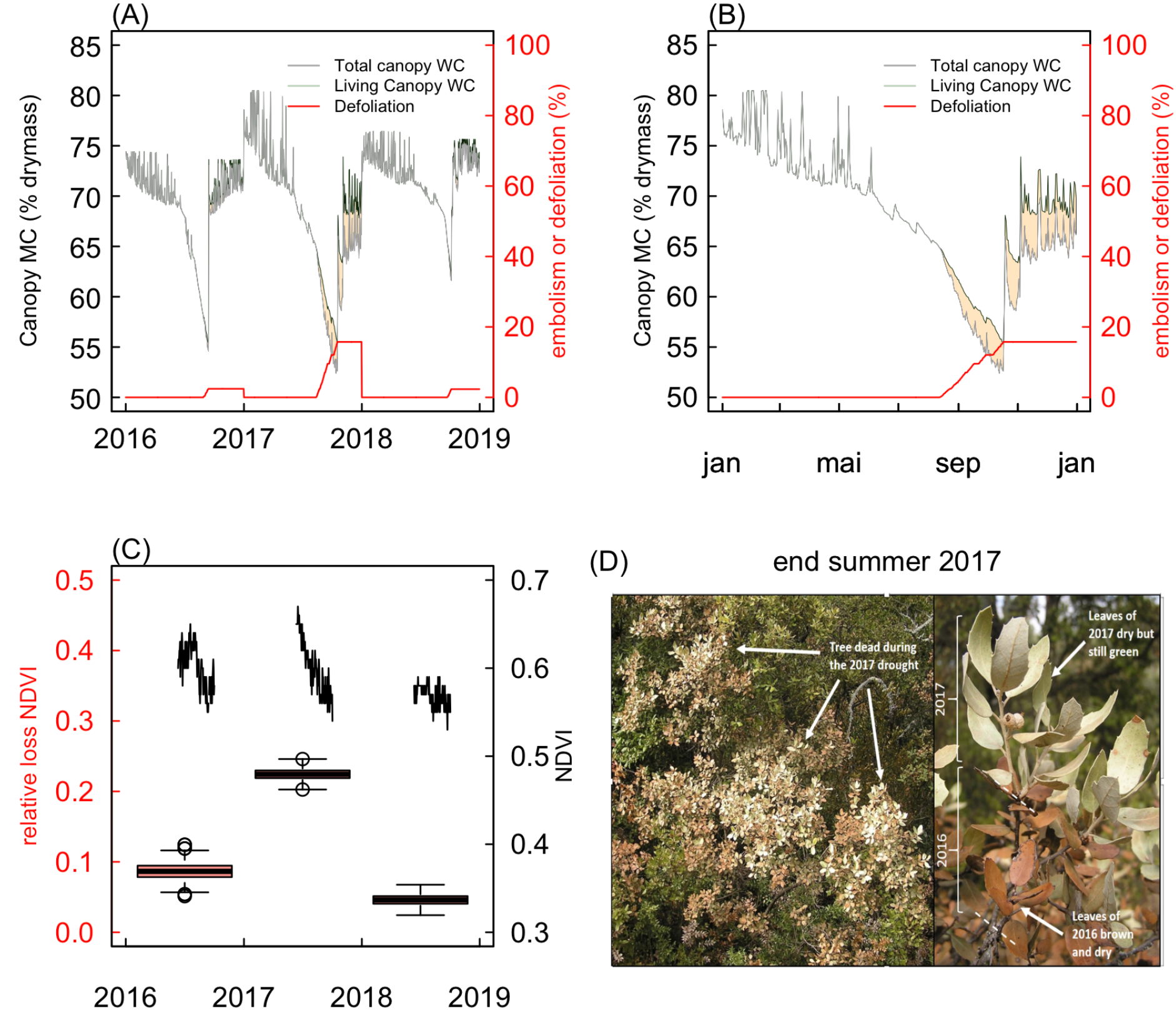
Simulated canopy level moisture content dynamics for the studied period (A and B) and measured indicators of foliage mortality (C, D). For the three studied years (panel A) and the driest year (panel B) the dynamic of the canopy moisture content of the living and dead fraction of the canopy are shown. The simulated foliage mortality is also indicated. The panel C shows NDVI dynamics during the vegetation season of the three studied years as well the standardized variation (boxplot) leaf maturity (early July) and the end of summer drought (mid-October). The panel D is a picture taken at the end of summer 2017 showing exceptional leaf desiccation occurring following the extreme drought. Note that every year the leaf area index and plant hydraulic conductance are reset to their initial value.

#### Sensitivity analysis to traits and climate change

The Figure 6 shows how an increase of some key model parameters by 40 % around the default value (Table 1), change model outputs in terms of minimal leaf level and canopy level moisture content and foliage mortality. The quantile 5 percent of leaf *FMC* and canopy *FMC* for daily simulations in the 1999-2019 period were used. At leaf level the most sensitive traits included those acting on the water balance, such the water potential at stomatal closure (*ψ_close_*), the leaf area index (*LAI*) and the soil water capacity (*SWC*), but also traits involved in the relationship between water potential and RWCs. In particular the osmotic potential at full turgor (π_0_), which determines the minimum value of *RWC*_symp_ for a given water potential has a large effect on *LFMC*. By contrast *P_50_*, *g_min_* and 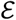 had relatively a low effect (<5%). For canopy level *LFMC* the water balance parameters had even a greater role and so it was for the *P*_50_. This was fully explained by the contribution of these traits to leaf mortality that was due to drought induced embolism in the model (Figure 5). Simulation under climate change showed an overall decreasing trend in *LFMC* at leaf and canopy scale with an increased in leaf mortality.

**Figure 6:**
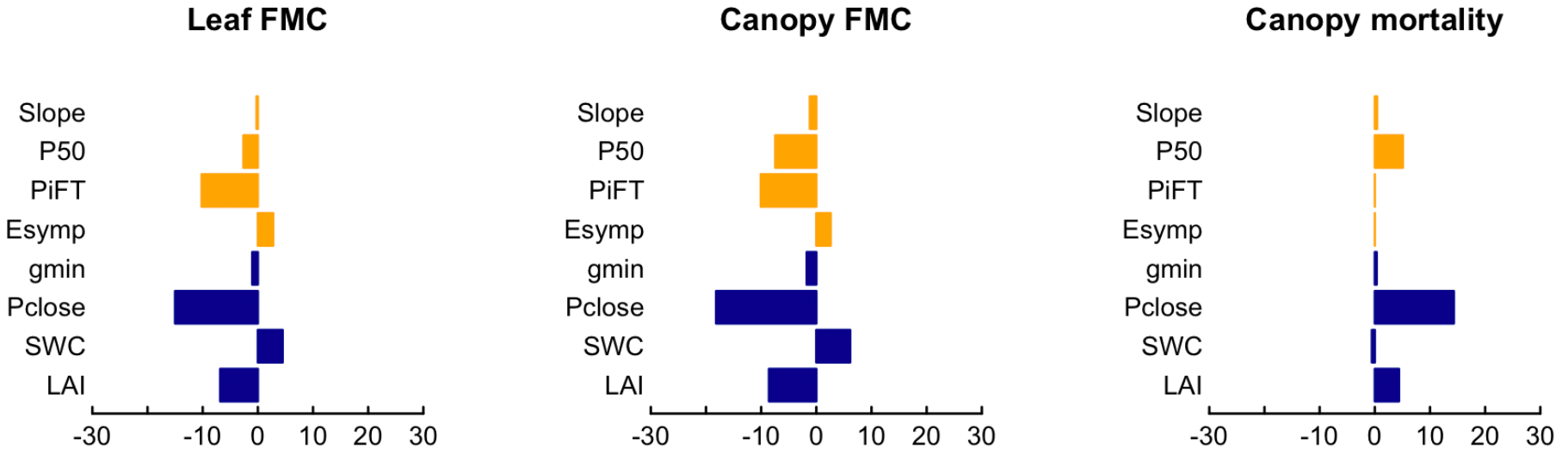
Sensitivity analysis of leaf and canopy fuel moisture content and foliage mortality to a 40% increase (from −20% to + 20% around the default parameters value, Table 1) of key model parameters. Physiological traits driving the relationship between leaf level moisture content or leaf mortality and water potential are colored in orange and parameters involved in the water balance are colored in blue. The parameters tested include the vulnerability curve to cavitation (slope and P50), the pressure volume curves (Osmotic potential at full turgor (PiFT and Esym), the minimal leaf conductance to water vapor (gmin), the water potential at stomatal closure (Pclose) and the available soil water capacity (SWC) and the leaf area index (LAI).

**Figure 7:**
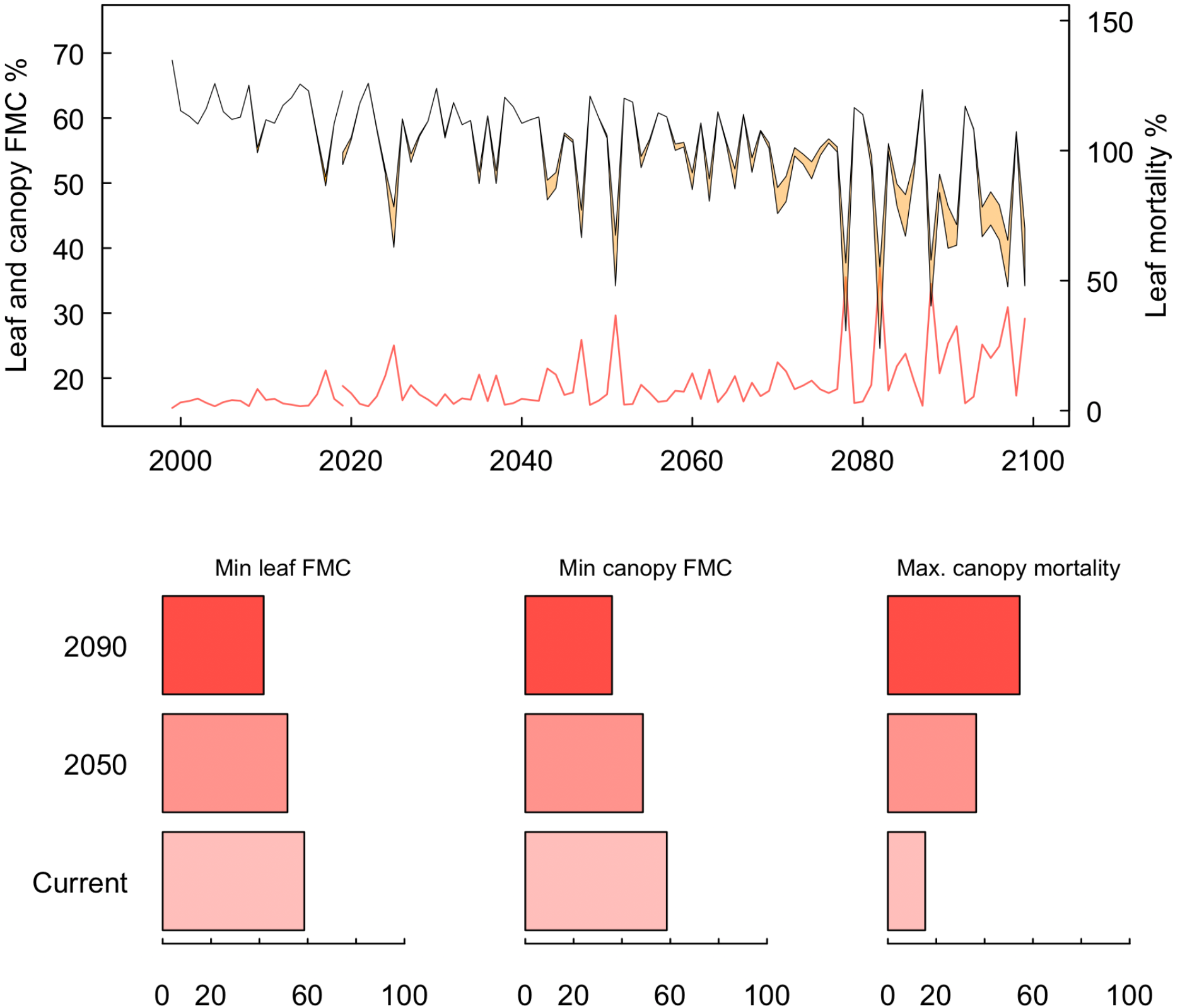
Modelled yearly minimum leaf and canopy fuel moisture content and maximum foliage mortality in response to climate change. Daily climate data derived from one coupled GCM-RCM model (MPI – ESM coupled with Remo2009) representative of the average of 5 different models in terms of drought (see Fargeon et al 2019). Only one climate scenario from CMIP 5 (RCP8.5). The is no interannual memory in these simulations and parameters value were reset to their default value (Table 1) at the beginning of each year in spite of potential decline due to drought (in particular leaf area index and plant hydraulic conductance).

## Discussion

Live fuel moisture content is a key trait involved in fire danger prevention which dynamics is understood or predicted. Empirical drought indices predict poorly LFMC variations (Soler Martin *et al.* 2017; Ruffault, Martin-StPaul, *et al.* 2018) and among physiological traits, water potential has been shown to be tightly correlated to LFMC dynamics (Nolan *et al.* 2018; Pivovaroff *et al.* 2019). However no studies have explored the mechanistic linkage between LFMC and water potential during drought. In the following section we described how our approach allowed to identify key traits involved in LFMC dynamics during drought at the leaf and canopy level.

### Leaf level LFMC dynamic depends on water potential osmotic adjustment

Leaf LFMC values recorded during three consecutive summer drought could be predicted from leaf water potential measurement using a set of mechanistic equations representing state of the art ecophysiological hydraulic processes (Figure 2). Two types of process were accounted: the pressure volume curves theory driving the symplasmic water content and leaf xylem vulnerability to cavitation for the apoplasmic part.

Most of the leaf level LFMC variations measured can be attributed to symplasmic variation for different reasons. First the symplasmic compartment is elastic and its water content decrease for relatively high water potential, consistent with the range measured *in situ* spanned from −1 to −5.5 MPa. By contrast, the apoplasmic compartment is made inelastic and empties for very negative water potential that causes cavitation. In our case, even the lowest values of water potential reached in 2017 caused less than 30% embolism. Indeed, *Quercus ilex* is highly resistant to embolism (Sergent *et al.* 2020). Leaf vulnerability measured for trees on the site yielded a *P*_50_ averaging at −6.1 (MPa+− 1.4MPa; Moreno et al in preparation; Martin-StPaul et al in preparation) leading to a level of embolism below 30% even during the 2017 extreme drought, which is consistent with native embolism estimation made in march 2018 (data not shown). In addition, there is slightly more symplasmic than apoplasmic mass in leaves (symplasmic fraction 0.56).

By using the traits measured at the laboratory, the predictability of LFMC from water potential measurements was relatively high (R2= 0.55, Figure 2) and consistent with previous studies (Nolan *et al.* 2018; Pivovaroff *et al.* 2019). Interestingly by optimizing a few key parameters of the pressure volume curves (π_0_, ε, *FDMC*), we found values very similar to those derived from pressure volume curves measured in 2018. This result indicates that LFMC can be predicted from water potential by using pressure volume curves parameters -- between dates along a dry season (and even between years)-- with the same accuracy as if the model was optimized with long term field measurements.

However, and in spite of this encouraging result, the model was significantly improved (Table 2) by assuming that osmotic adjustment (i.e. change in the value of π_0_) occurs from one year to the other (Figure 2, Table 2). Such adjustment changes the relationship between bulk water potential and water content (Figure 2). Osmotic adjustment is known to be an acclimation mechanism allowing to maintain tissue hydration and turgor in spite of water potential decrease and that occurs for many species (Bartlett *et al.* 2012), including *Quercus ilex* (Limousin et al in prep). Such process complicates the predictability of leaf LFMC as there is so far no generic model including osmotic adjustment (but see (Rieger 1995)). Future experimental research should measure concurrently water potential, water content and osmotic potential to improve our understanding of their inter-relation.

### Canopy level LFMC depends on hydraulic failure and leaf mortality

Depending on the scientific community, LFMC is assessed at the leafy shoot level –this is the case for field *in situ* measurements (Martin-StPaul *et al.* 2018; Yebra *et al.* 2019) -- or at the scale of stand canopy which is the case for remote sensing measurements (Yebra *et al.* 2013). The model presented here attempt to bridge the gap between both scales by integrating leaf level estimates of LFMC at the canopy level and by accounting for leaf mortality during extreme drought. Accordingly, simulations along the studied period showed that most of the time, canopy LFMC is dictated by leaf level variations (Figure 4) as leaf mortality rarely occurs. However, during the extreme drought 2017 foliage mortality reached almost 20 % and significantly contributed to canopy level LFMC decline (Figure 5).

The detailed mechanisms involved in drought-induced mortality of leaves are so far poorly known. Here we assumed that drought induced xylem embolism lead to a proportional disconnection between the leaves and the plant, thereby driving leaf mortality. This hypothesis appears parsimonious as it does not rely on additional processes than those already accounted for and is corroborated by various experimental data (Barigah *et al.* 2013; Urli *et al.* 2013). Through this assumption, we simulated a leaf mortality peak during the extreme drought 2017 which is consistent with (i) the level of embolism recorded on the site in march 2018, (ii) NDVI decline that occurred during this specific drought but not during previous and following years when drought intensity was lower (Figure 5) and (iii) *in situ* observation of leaf browning (Figure 5b). However, it must be acknowledge that active processes might also be involved in leaf mortality and desiccation which could shift the pattern of leaf desiccation from those of hydraulic failure in other situation (e.g. other species). More research and experimental data on leaf shedding are necessary to improve understanding and predictability of canopy LFMC under extreme drought.

### Water balance and hydraulic parameters drives the LFMC dynamic during drought

The sensitivity analysis revealed the differential role of some key parameters on the LFMC dynamics at leaf of canopy scale during drought. First of all, parameters affecting the water balance (i.e. the level of transpiration before or during the drought), and thereby the water potential decline have an important influence on both leaf and canopy level LFMC dynamic. Indeed, water potential declines directly affect water content through the physiological relationship, but also leaf mortality through drought induced embolism. In particular, *ψ_close_* (the water potential causing stomatal closure) had the greatest effect on minimum LFMC values. As discussed in a previous study, the high non linearity in the pedo-transfer function is responsible for this pattern (Martin-StPaul et al 2017). Transpiration has an increasingly important effect on water potential – and thus on LFMC decline – as water potential is becoming more negative. Hence, a later stomatal closure implies a greater fuel desiccation. Then leaf area index and soil water capacity, that affect directly the water balance and thus the water potential have an effect of the order of 5 to 10% on minimum leaf LFMC values. However *g*_min_ has a relatively small effect in the range considered here (from 2.4 to 3.6). Secondly, traits driving the relationship between water potential and moisture content have different sensitivity between leaf and canopy level. The pressure volume curve parameters (in particular the π_0_) presented the highest effect at the leaf level compared to the vulnerability to cavitation parameters (in particular the slope). Interestingly, this difference reduced considerably at the canopy scale as *P*_50_ is the main drivers of canopy mortality and canopy desiccation. Our simulations show that under the climate conditions projected to 2100, the contribution of foliage mortality in determining canopy LFMC increases considerably (Figure 6). It has to be noted that osmotic adjustment was not considered in these simulations, but this would have reinforced the contribution of canopy mortality to canopy LFMC as osmotic adjustment increases water content at the leaf level.

### Summary, perspectives and conclusions

Our analysis allowed to identify some key mechanisms responsible for LFMC dynamics during extreme drought at the leaf and the canopy level. Such analysis should be extended on multiple species and could be improved by some way. For instance, we did not considered the drivers of fuel dry matter content in this approach, assuming that this process can be neglected during extreme drought. This is likely to be reasonable for current year leaves, that are produced mostly before the drought period for this species. However, LDMC variations are likely to be influent at the interannual scale when the drought period overlap with the leaf and shoot growth. In the future, extending the approach to multiple species should consider phenology and an evaluation of the effects of water status on dry matter accumulation to improve the predictability of leaf level dynamics of LFMC (Jolly *et al.* 2014).

Our study, also show that the parameters driving stand water balance appears crucial to determine water potential and thus LFMC dynamics under drought. To produce large scale predictions, some of these parameters could be informed using global data base (for soil available water capacity for instance, *ψ_close_* or *g*_min_) or remote sensing products (for leaf area index). We did not test the model sensitivity to soil texture, despite we are aware it could have an important effect (Sperry *et al.* 1998; Martin-StPaul *et al.* 2017) that deserved to be studied also experimentally. Regarding the parameters affecting the plant desiccation at a given water potential, it appears that pressure volume curve parameters have a dominant effect under “normal” climatic condition, which makes it possible to explore LFMC for multiple species. However, osmotic adjustment have a great influence at leaf level and is known to occur on multiple species (Bartlett *et al.* 2012). Its drivers should be explore on multiple species.

Under future conditions, vulnerability to cavitation seems to have an increasingly great importance in that sense that it drives foliage mortality. Here again, research efforts are needed to better determine whether foliage mortality occurs concurrently of before drought induced embolism so that it would be possible to extend the modelling approach to multiple species. The segmentation of the vulnerability to cavitation trees appears in general modest for trees (Li *et al.* 2020), however, active processes could drive leaf shedding in some species as a mean to save water and dampen drought effects over the long run (Choat *et al.* 2018).

## Notes

### Competing Interest Statement

The authors have declared no competing interest.

